# Plasticity in airway smooth muscle differentiation during mouse lung development

**DOI:** 10.1101/2022.05.27.493761

**Authors:** Katharine Goodwin, Bezia Lemma, Adam Boukind, Celeste M. Nelson

## Abstract

Smooth muscle differentiation has been proposed to sculpt airway epithelial branches in mammalian lungs. Serum response factor (SRF) acts with its cofactor myocardin to promote the expression of contractile smooth muscle markers. However, smooth muscle cells exhibit a variety of phenotypes beyond contractile that are independent of SRF/myocardin-induced transcription. To determine whether airway smooth muscle exhibits phenotypic plasticity during embryonic development, we deleted *Srf* from the pulmonary mesenchyme. *Srf*-mutant lungs branch normally, and the mesenchyme exhibits normal cytoskeletal features and patterning. scRNA-seq revealed an *Srf*-null smooth muscle cluster wrapping the airways of mutant lungs that lacks contractile smooth muscle markers but retains many features of control smooth muscle. *Srf-*null airway smooth muscle exhibits a synthetic phenotype, compared to the contractile phenotype of mature wildtype airway smooth muscle. Our findings reveal plasticity in airway smooth muscle differentiation and demonstrate that a synthetic smooth muscle layer is sufficient for airway branching morphogenesis.

## Introduction

Vascular and visceral smooth muscle cells are present throughout the body and their differentiation is key for development and homeostasis (Jaslove and Nelson, 2018, Donadon and Santoro, 2021). In the adult, smooth muscle can take on contractile or synthetic/proliferative phenotypes in response to local signals (Owens et al., 2004, Rensen et al., 2007, Wang et al., 2015). For vascular smooth muscle, scRNA-seq analysis has revealed at least six phenotypes (Yap et al., 2021). Studies using cultured cells have implicated myocardin, acting with its required co-factor, serum response factor (SRF), as a key transcriptional activator of the expression of contractile smooth muscle markers (Du et al., 2003, Wang et al., 2003), including *Acta2* (α-smooth muscle actin; αSMA), *Tagln* (transgelin, SM22α), and *Cnn1* (calponin-1). Changes in *Myocd* expression are associated with cardiovascular disease, and downregulation of *Myocd* can lead to phenotypic switching of vascular smooth muscle to an osteogenic phenotype and subsequent calcification of arteries in patients with type 2 diabetes (Miano, 2015).

SRF-deficient embryos die at embryonic day 6.5 (*E*6.5) from a defect in gastrulation due to a complete loss of mesenchymal cells (Arsenian et al., 1998). Myocardin (*Myocd*) -/-animals die at *E*10.5 as a result of vascular abnormalities (Li et al., 2003). Injecting *Myocd*-/-embryonic stem cells into wildtype blastocysts revealed that wildtype cells outcompete myocardin-null cells in visceral smooth muscle, but less so in vascular smooth muscle (Hoofnagle et al., 2011). SRF/myocardin signaling was therefore concluded to be essential for visceral smooth muscle differentiation. Nonetheless, mice deficient in *Acta2, Tagln*, or *Cnn1* are all viable (Zhang et al., 2001, Feng et al., 2019, Schildmeyer et al., 2000), suggesting that these SRF/myocardin targets are not required for smooth muscle differentiation.

The airway smooth muscle differentiation program is just beginning to be defined. Multiple signaling pathways affect each step of the differentiation program, from regulating the pool of smooth muscle progenitors, to specifying nascent smooth muscle cells, to inducing their differentiation and maturation (Goodwin et al., 2022). Importantly, cytoskeletal and adhesion genes are expressed before contractile smooth muscle markers, suggesting early changes to cell mechanical properties as the mesenchyme begins to differentiate into smooth muscle.

Disrupting the smooth muscle differentiation program leads to defective branching morphogenesis of the epithelium, with a loss of smooth muscle differentiation associated with cystic branches and a gain associated with stunted branches (He et al., 2017, Boucherat et al., 2015, Yi et al., 2009, Goodwin et al., 2019, Goodwin et al., 2022, Jaslove et al., 2022, Kim et al., 2015). These observations have led to a conceptual model wherein the pulmonary mesenchyme sculpts the morphology and positions of branches that form in the growing airway epithelium (Jaslove and Nelson, 2018).

However, recent work in which *Myocd* was deleted from the embryonic pulmonary mesenchyme appears to cast doubt on this model. *Myocd*-mutant lungs fail to express markers of contractile smooth muscle but branch normally (Young et al., 2020). That study did not determine whether loss of Myocd affects the entire smooth muscle differentiation program, raising the possibilities that either the early stages of smooth muscle differentiation (which sculpt new epithelial branches and occur prior to robust *Myocd* expression) are unaffected by loss of *Myocd*, and/or that *Myocd*-independent aspects of the smooth muscle gene expression program are sufficient to support epithelial morphogenesis. Here, we sought to resolve these seemingly contradictory observations and define precisely the role of SRF/myocardin-induced transcription in the embryonic pulmonary mesenchyme.

## Results

### Deleting Srf from the embryonic pulmonary mesenchyme permits airway branching but decreases airway diameter

To understand the role of SRF/myocardin-induced transcription in airway smooth muscle differentiation and morphogenesis in the embryonic mouse lung, we deleted *Srf* from the pulmonary mesenchyme using *Tbx4-rtTA;tet-O-Cre* (Zhang et al., 2013). This strategy allows for doxycycline-induced activation of Cre recombinase exclusively in the pulmonary mesenchyme by using a tissue-specific enhancer of *Tbx4*. Immunofluorescence analysis for E-cadherin (Ecad) and α-smooth muscle actin (αSMA) revealed that deleting *Srf* from the pulmonary mesenchyme results in a normally branched airway epithelial tree devoid of αSMA^+^ cells at *E*12.5 (**Fig. 1A-B**). The number of terminal branches is similar in *Srf*-mutant and littermate control lungs (**Fig. 1C**), and we observed no differences in lung size based on projected area of the epithelium (**Fig. S1A-B**) or the whole lung (**Fig. S1C-D**). However, *Srf*-mutant lungs have significantly smaller caliber airways based on measurements of the left primary bronchus between the first and second domain branches (**Fig. 1D**).

**Figure 1.**
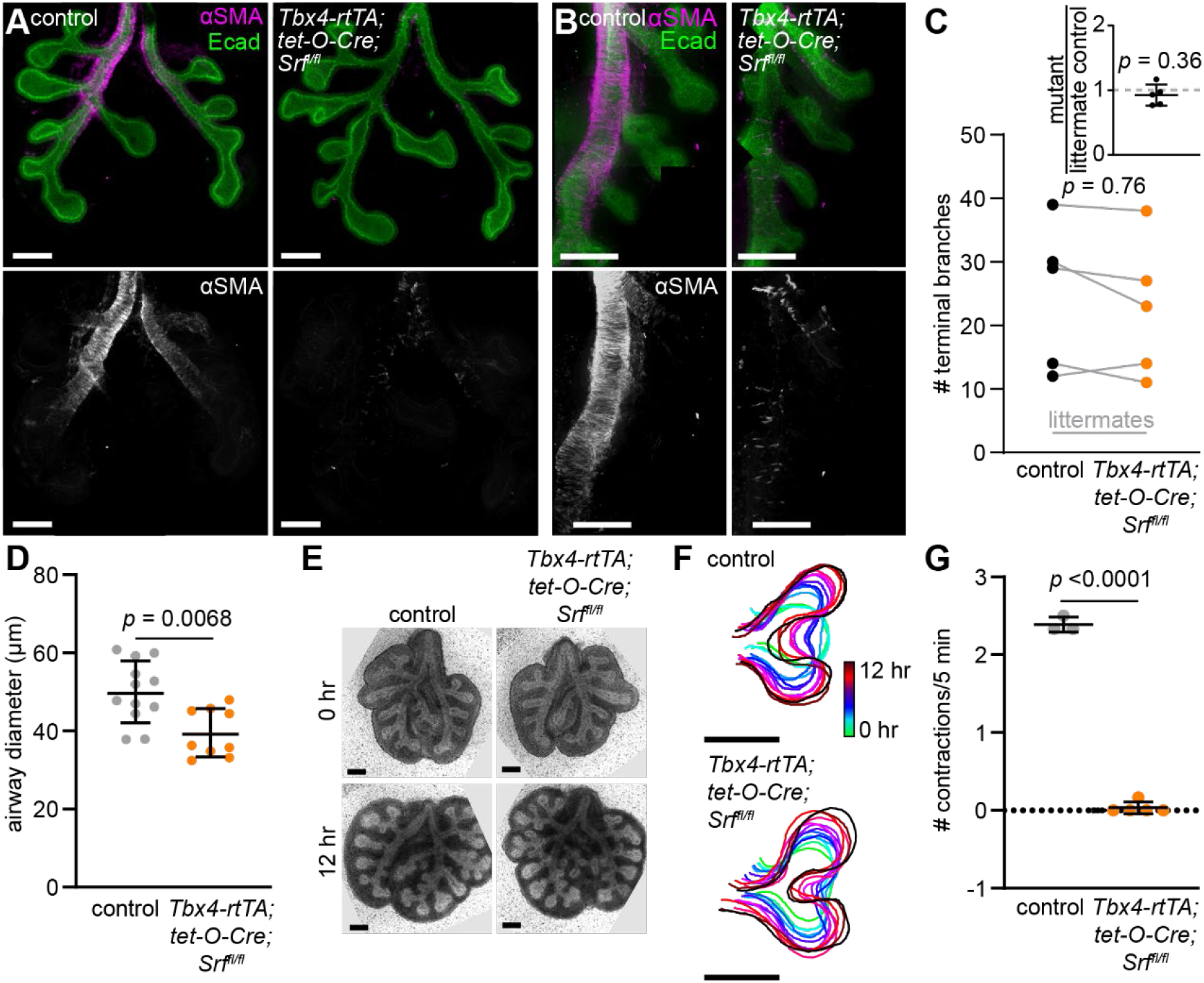
Deleting *Srf* from the embryonic pulmonary mesenchyme leads to loss of αSMA but has no effect on epithelial branching morphogenesis. (**A-B**) *E*12.5 *Tbx4-rtTA;tet-O-Cre;Srf*^*fl/fl*^ and littermate control lungs immunostained for Ecad and αSMA. Scale bars indicate 100 μm. (**C**) Number of branches in *E*12.5 *Tbx4-rtTA;tet-O-Cre;Srf*^*fl/fl*^ and littermate controls, connected by gray lines and compared using a t-test. Inset shows branch number in mutant lungs divided by that of littermate controls compared to a hypothetical mean of 1 using a one sample t-test. Error bars show s.d. (**D**) Airway diameter measured between the first two domain branches of the left lobe of *E*12.5 *Tbx4-rtTA;tet-O-Cre;Srf*^*fl/fl*^ lungs and controls (n = 12 controls, 9 mutants). (**E**) Brightfield images of *E*12.5 *Tbx4-rtTA;tet-O-Cre;Srf*^*fl/fl*^ and littermate control lungs cultured *ex vivo*. Scale bars indicate 200 μm. (**F**) Contours of the epithelium showing the dynamic changes in the morphology of terminal bifurcations in mutants and controls. Scale bars indicate 50 μm. (**G**) Number of spontaneous smooth muscle contractions (assessed by airway deformation) per five-minute timelapse in *Tbx4-rtTA;tet-O-Cre;Srf*^*fl/fl*^ and littermate control lungs (n = 3 controls, 5 mutants). Error bars show s.d. Groups were compared using two-sided t-test.

We used *ex vivo* culture and live-imaging of *Srf*-mutant lungs and controls to visualize the dynamics of branching morphogenesis and airway smooth muscle contractions (**Fig. 1E**). Bifurcation of the airway epithelium proceeds at the same rate in *Srf* mutants as in controls (**Fig. 1F**), consistent with their ability to form a normal number of branches. However, *Srf*-mutant airways lack spontaneous contractile behaviors (**Fig. 1G**). These observations are similar to previous descriptions of lungs from mice in which the SRF cofactor *Myocd* had been deleted specifically from the embryonic pulmonary mesenchyme (Young et al., 2020). Combined with that work, our data suggest that SRF/myocardin-dependent genes are not required in the pulmonary mesenchyme for airway epithelial branching.

### Srf-mutant lungs have normal mesenchymal patterning and mechanical properties

Given that deleting *Srf* from the mesenchyme has no effect on epithelial morphogenesis, we next focused our attention on the mesenchyme itself to characterize the phenotype of this mutant. Specifically, we evaluated the patterning and mechanical properties of the mesenchyme in *E*12.5 *Srf*-mutant lungs. In wildtype lungs, the branching epithelium is surrounded by sub-epithelial mesenchymal cells that express lymphoid enhancer binding factor 1 (Lef1) and bone morphogenetic protein 4 (Bmp4) (**Fig. 2A**) (Goodwin et al., 2022). Immunofluorescence and RNAscope analysis revealed that Lef1 and Bmp4 are expressed in similar patterns in *Srf* mutants and controls (**Fig. 2B-D**), indicating that *Srf* is not required for patterning of the sub-epithelial pulmonary mesenchyme. In wildtype lungs, nascent smooth muscle cells immediately adjacent to the epithelium express T-box transcription factor 3 (Tbx3) (Goodwin et al., 2022). Immunofluorescence analysis revealed that Tbx3 is also expressed in the mesenchyme adjacent to the epithelium in *Srf* mutants (**Fig. 2E-F**), suggesting that Srf is similarly dispensable for specifying the earliest stages of the airway smooth muscle differentiation program.

**Figure 2.**
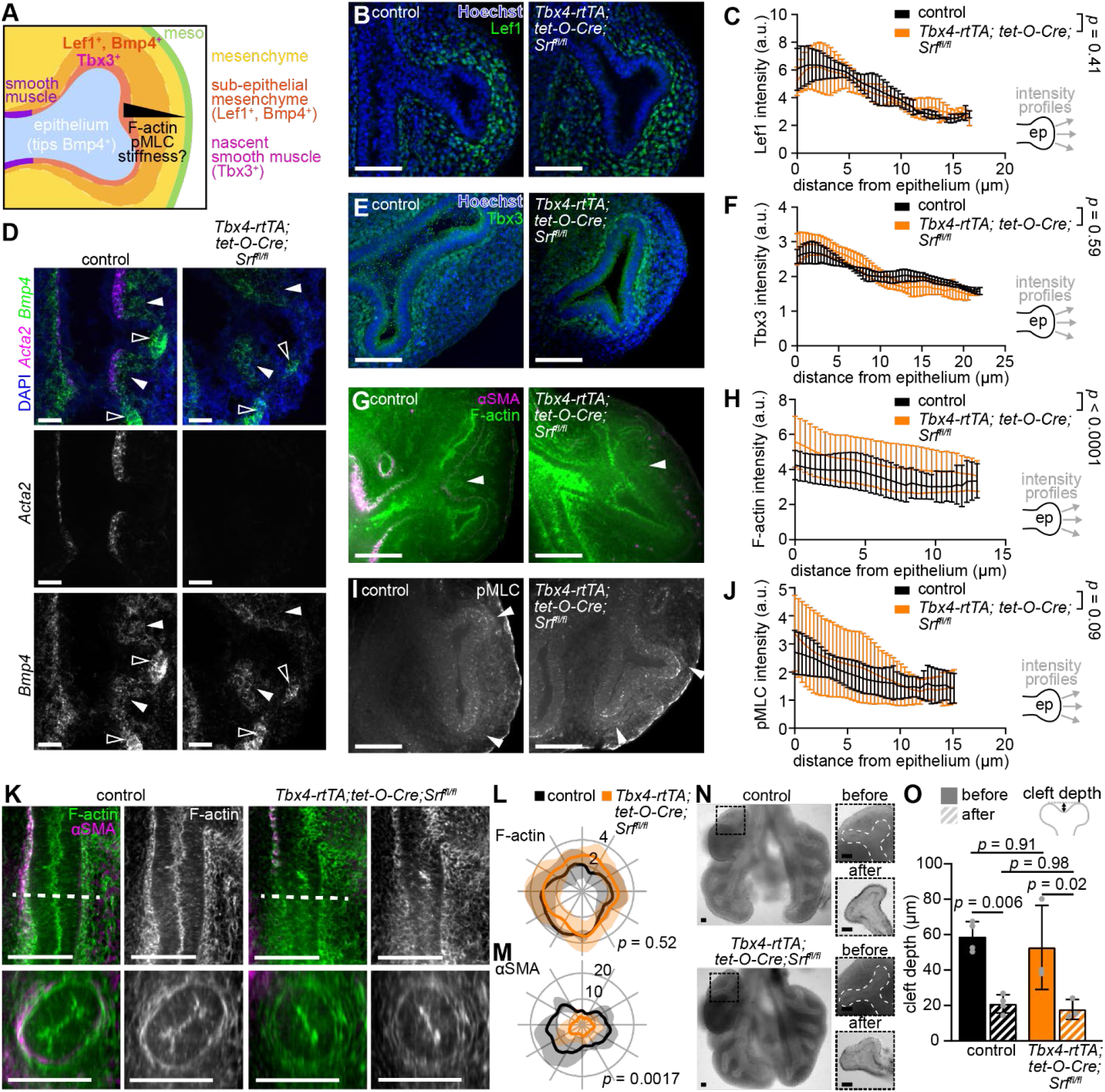
*Srf-*mutant lungs exhibit normal mesenchymal patterning. (**A**) Schematic illustrating the expected patterns of cytoskeletal elements, tissue mechanical properties, and markers of patterning in the mesenchyme around epithelial tips. (**B-C**) *E*12.5 control and *Tbx4-rtTA;tet-O-Cre;Srf*^*fl/fl*^ lungs immunostained for Lef1 and counterstained with Hoechst and quantification of Lef1 intensity profiles around epithelial buds (n = 3 controls, 3 mutants). Mean and s.d. are plotted and curves were compared using two-way ANOVA. Schematic shows lines and direction along which intensity profiles were measured. (**D**) Fluorescence *in situ* hybridization for *Acta2* and *Bmp4* on sections of *E*12.5 control and *Tbx4-rtTA;tet-O-Cre;Srf*^*fl/fl*^ lungs. Filled arrowheads indicate *Bmp4*^*+*^ sub-epithelial mesenchyme and empty arrowheads indicate *Bmp4*^*+*^ epithelial tips. (**E-F**) *E*12.5 control and *Tbx4-rtTA;tet-O-Cre;Srf*^*fl/fl*^ lungs immunostained for Tbx3 and counterstained with Hoechst and quantification of Tbx3 intensity profiles around epithelial tips (n = 3 controls, 3 mutants). Mean and s.d. are plotted and curves were compared using two-way ANOVA. (**G-J**) *E*12.5 control and *Tbx4-rtTA;tet-O-Cre;Srf*^*fl/fl*^ lungs immunostained for αSMA and F-actin or pMLC and quantification of F-actin and pMLC intensity profiles around epithelial tips (n = 9 controls, 11 mutants for F-actin; n = 4 controls, 6 mutants for pMLC). Arrowhead indicates enrichment of mesenchymal F-actin and pMLC near the branching epithelium. (**K**) Single confocal slices and reconstructed optical cross-sections through *E*12.5 control and *Tbx4-rtTA;tet-O-Cre;Srf*^*fl/fl*^ lungs immunostained for αSMA and F-actin. The dashed line indicates the location of the cross-section. (**L-M**) Polar plots showing F-actin and αSMA intensity around the airways based on angle from the center of the bronchus in *E*12.5 control and *Tbx4-rtTA;tet-O-Cre;Srf*^*fl/fl*^ lungs (n = 3 controls, 9 mutants). Line shows mean and shaded area shows s.d. *p* value is for the comparison of mean intensities using two-sided t-test. (**N**) Brightfield images of *E*12.5 control and *Tbx4-rtTA;tet-O-Cre;Srf*^*fl/fl*^ lungs, with zoomed-in regions showing bifurcations before and after surgical removal of the mesenchyme. (**O**) Quantification of the depths of epithelial clefts in mutants and controls before and after surgical removal of the mesenchyme (n = 4 controls, 3 mutants). Scale bars indicate 50 μm.

Filamentous actin (F-actin) and phosphorylated myosin light chain (pMLC) are molecular markers that are associated with the mechanical properties of airway smooth muscle – specifically, enhanced stiffness and mechanical tone (Sieck et al., 2019, Luo et al., 2019, Gunst and Zhang, 2008). Consistently, the levels of F-actin and pMLC are enriched in the mesenchyme immediately adjacent to epithelial tips, sites of active branching morphogenesis, and decrease farther from the epithelium (**Fig. 2G-J**). We observed similar patterns of F-actin and pMLC in *Srf* mutants (**Fig. 2G-J**), with slightly higher levels of F-actin in the mesenchyme of *Srf*-mutant lungs (**Fig. 2H**). Furthermore, the highest levels of F-actin are concentrated in the mesenchyme wrapping around epithelial branches in both controls and mutants (**Fig. 2K-L**), despite loss of αSMA in the latter (**Fig. 2M**). These data suggest that loss of Srf does not decrease the stiffness or mechanical tone of the pulmonary mesenchyme.

Based on the normal patterning of these molecular markers of mechanical stiffness and tone, we hypothesized that the *Srf*-null mesenchyme remains capable of mechanically sculpting the airway epithelium. To test this hypothesis, we compared the morphology of epithelial bifurcations before and after surgically removing the surrounding mesenchyme. In both mutants and controls, the depth of the cleft at bifurcations significantly decreases following removal of the mesenchyme (**Fig. 2N-O**). Importantly, cleft depths in *Srf* mutants are initially indistinguishable from those in controls, and they decrease to identical levels after removing the mesenchyme.

These results suggest that the forces exerted by the mesenchyme on the epithelium are indistinguishable in *Srf* mutants and controls. Based on these data, we conclude that *Srf* is not required for patterning the molecular or passive mechanical features of the embryonic pulmonary mesenchyme. As a consequence, mesenchymal sculpting of the epithelium is independent of *Srf*-induced transcription.

### scRNA-seq of Srf-mutant lungs reveals a “smooth muscle-like” population of cells

Given that deleting *Srf* does not alter known mesenchymal patterning around the airway epithelium, we used single-cell transcriptomics to broadly characterize the changes induced by loss of *Srf*. We generated scRNA-seq datasets from whole *Tbx4-rtTA;tet-O-Cre;Srf*^*fl/fl*^ lungs and *Srf*^*fl/fl*^ controls isolated at *E*12.5 (**Fig. 3A**). Both datasets contained the expected populations of cells, including mesenchyme, epithelium, vascular endothelium, mesothelium, neurons, and immune cells (**Fig. 3A and Fig. S2A**). Consistent with our immunofluorescence analysis, the SRF-induced contractile smooth muscle markers *Acta2* and *Tagln* were detected in the smooth muscle cluster of the control dataset (cluster 7) and were absent from the mutant dataset (**Fig. 3B**). *Srf* was detected throughout the control mesenchyme and largely absent from the mutant mesenchyme, confirming that mesenchymal expression of Cre resulted in the expected knockout (**Fig. 3C**). *Myocd* was present in the smooth muscle cluster in the control dataset, as expected, but was also expressed at high levels in cluster 9 of the mutant dataset (**Fig. 3C**). As compared to the smooth muscle cluster in the control, cells in the mutant cluster 9 expressed significantly lower levels of canonical markers of contractile smooth muscle, including the aforementioned *Acta2* and *Tagln*, as well as *Cnn1, Myl9*, and *Myh11* (**Fig. S2B-F**). This *Myocd*^+^ cluster also did not contain *Srf*-expressing cells (**Fig. 3C, Fig. S2G**), ruling out the possibility of a failure in Cre-mediated recombination.

**Figure 3.**
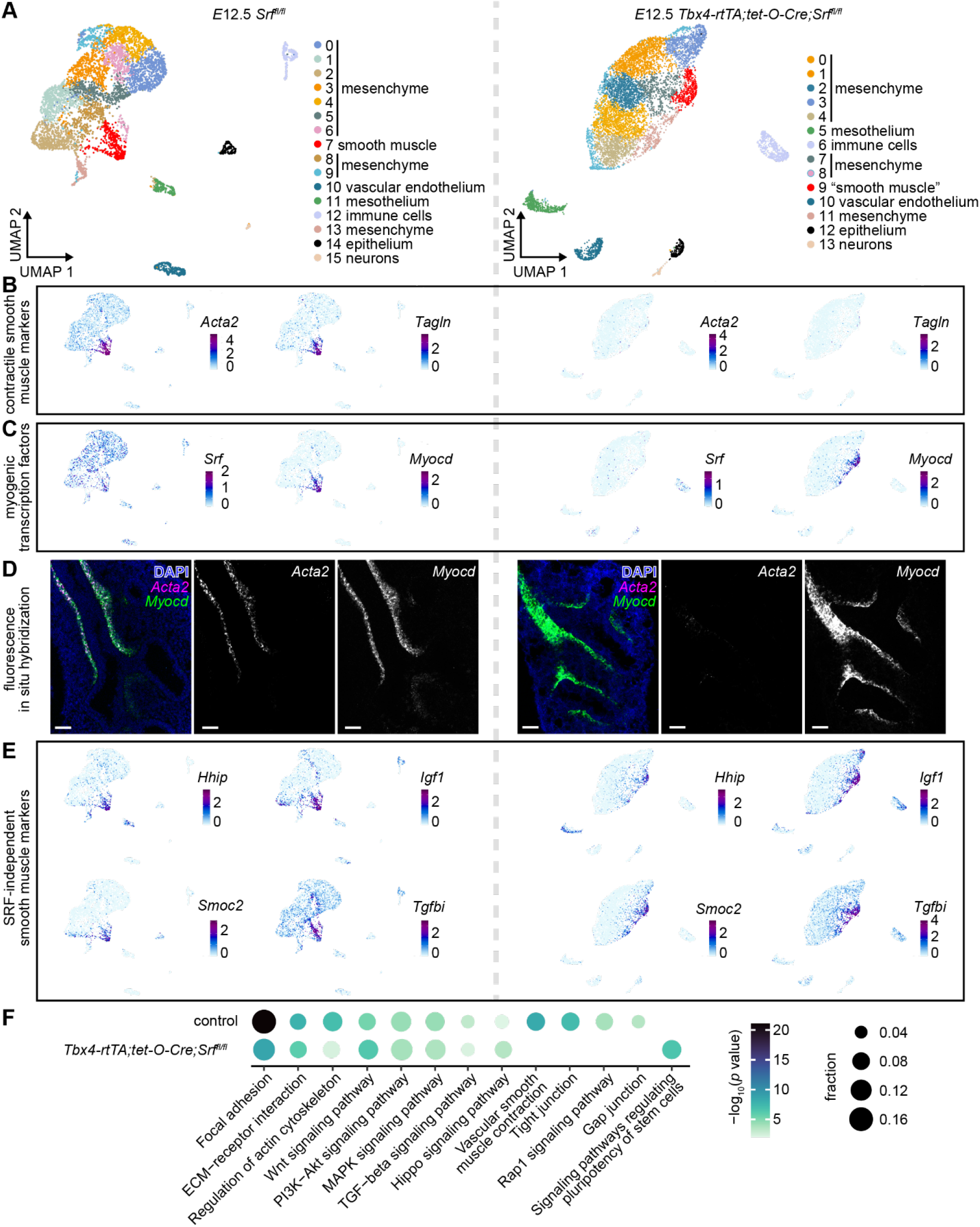
scRNA-seq reveals a smooth muscle-like population in *Srf*-mutant lungs. (**A**) UMAPs of cells isolated from *E*12.5 *Tbx4-rtTA;tet-O-Cre;Srf*^*fl/fl*^ and littermate control lungs color-coded according to cluster (n = 6 controls, 7 mutants). (**B**) Expression levels of contractile smooth muscle markers *Acta2* and *Tagln* in mutants and controls. (**C**) Expression levels of myogenic transcription factors *Srf* and *Myocd* in mutants and controls. (**D**) Fluorescence in situ hybridization for *Acta2* and *Myocd* in *E*12.5 *Tbx4-rtTA;tet-O-Cre;Srf*^*fl/fl*^ and littermate control lungs. Scale bars indicate 50 μm. (**E**) Expression levels of SRF-independent smooth muscle markers *Hhip, Igf1, Smoc2*, and *Tgfbi* in mutants and controls. (**F**) KEGG pathway enrichment analysis for marker genes of the smooth muscle clusters from control and *Tbx4-rtTA;tet-O-Cre;Srf*^*fl/fl*^ lungs.

To map the locations of the *Myocd*^+^ clusters, we carried out fluorescence *in situ* hybridization analysis of *Acta2* and *Myocd* transcripts in *E*12.5 control and mutant lungs. In control lungs, *Acta2* and *Myocd* expression showed close spatial overlap in cells adjacent to the epithelium, as expected for markers of airway smooth muscle (**Fig. 3D**). In contrast, *Srf*-mutant lungs lacked *Acta2* expression, in agreement with our immunofluorescence and scRNA-seq analyses. Nonetheless, the mutants exhibited clear *Myocd* expression in exactly the same pattern as controls. *Srf-*mutant lungs therefore contain a population of *Myocd*^+^ cells that wraps the airways in a pattern indistinguishable from that of airway smooth muscle. These data suggest that a “smooth muscle” cluster persists in lungs lacking mesenchymal *Srf*. Consistent with this conclusion, our scRNA-seq data revealed that many *Srf*-independent smooth muscle markers are expressed in the smooth muscle clusters of both control and *Srf*-mutant lungs (**Fig. 3E**).

To define the extent to which *Srf*-null smooth muscle is functionally similar to control smooth muscle, we carried out KEGG pathway enrichment analysis on marker genes for each smooth muscle cluster. Signaling pathways related to cell mechanical properties (e.g., focal adhesion, regulation of actin cytoskeleton) and to cell signaling (e.g., Wnt signaling, TGF-β signaling) were similarly enriched in both control and *Srf*-null smooth muscle (**Fig. 3F, Fig. S3**). Among the pathways unique to control smooth muscle were “vascular smooth muscle contraction” and “gap junctions”, suggesting that smooth muscle maturation may be impaired in *Srf* mutants (**Fig. 3F, Fig. S3**), in agreement with the loss of spontaneous contractions (**Fig. 1G**). Consistently, we found that genes under the KEGG term “signaling pathways regulating the pluripotency of stem cells” were enriched in only the mutant smooth muscle cluster (**Fig. 3F, Fig. S3**). These data reveal that deleting *Srf* from the embryonic pulmonary mesenchyme is not sufficient to eliminate airway smooth muscle. Instead, SRF-induced signaling appears to affect a minor subset of genes, including contractile smooth muscle markers.

### Srf-null smooth muscle cells exhibit a synthetic smooth muscle phenotype

Based on the above analysis, we hypothesized that the absence of either *Srf* or *Myocd* generates a population of mesenchymal cells encircling the airways that are transcriptionally and functionally similar to wildtype airway smooth muscle. To test this hypothesis, we asked whether *Srf*-null smooth muscle cells resemble nascent (immature) control smooth muscle, or whether they differentiate down a different trajectory. We carried out diffusion analysis (Haghverdi et al., 2015) of control mesenchymal cells, which revealed that the trajectory of undifferentiated mesenchyme bifurcates into either smooth muscle or cartilage precursors (**Fig. 4A, left**). We then projected the *Srf*-null smooth muscle cells onto this diffusion map (**Fig. 4A, right**). Mutant smooth muscle cells clustered among the undifferentiated mesenchymal cells near the bifurcation point between the smooth muscle and cartilage precursor branches. This position suggests that *Srf*-null smooth muscle may be trapped in the earliest stages of smooth muscle differentiation and prevented from maturing into a contractile phenotype.

**Figure 4.**
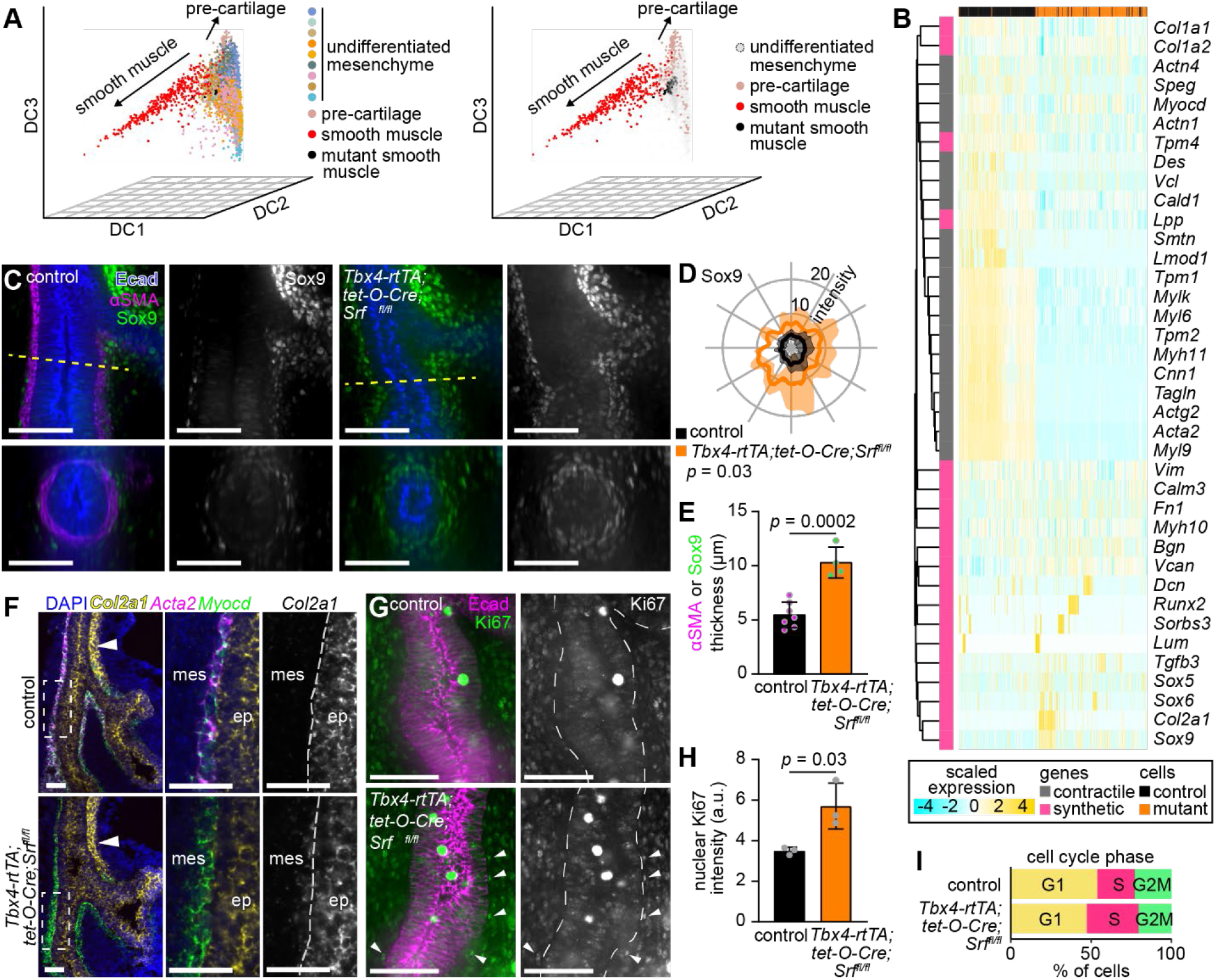
*Srf*-null airway smooth muscle cells display a synthetic smooth muscle phenotype. (**A**) Diffusion analysis of mesenchymal cells from the control dataset and projection of *Tbx4-rtTA;tet-O-Cre;Srf*^*fl/fl*^ mutant smooth muscle cells onto the diffusion map. Left plot is color-coded according to clusters from Fig. 3A; right plot is color-coded to emphasize control and mutant smooth muscle and cartilage precursors. (**B**) Clustering of mutant and control smooth muscle cells based on the expression of genes associated with contractile or synthetic smooth muscle phenotype. (**C**) Single confocal slices and reconstructed optical cross-sections through *E*12.5 control and *Tbx4-rtTA;tet-O-Cre;Srf*^*fl/fl*^ lungs immunostained for Ecad, αSMA, and Sox9. The dashed line indicates the location of the reconstructed cross-section. (**D**) Polar plots showing Sox9 intensity around the airways based on angle from the center of the bronchus of *E*12.5 control and *Tbx4-rtTA;tet-O-Cre;Srf*^*fl/fl*^ lungs (n = 5 controls, 2 mutants). Line shows mean and shaded area shows s.d. *p* value is for the comparison of mean intensities using two-sided t-test. (**E**) Thickness of the αSMA^+^ cell layer in controls and the Sox9^+^ cell layer in mutants compared by t-test (n = 7 controls, 4 mutants). (**F**) Fluorescence *in situ* hybridization for *Acta2, Myocd*, and *Col2a1* on sections of *E*12.5 control and *Tbx4-rtTA;tet-O-Cre;Srf*^*fl/fl*^ lungs. Arrowheads indicate *Col2a1*^*+*^ cartilage precursors along lateral side of the primary bronchus. Dashed boxes indicate zoomed-in regions shown to the right. Gray dashed line indicates border between the mesenchyme (mes) and epithelium (ep). (**G**) Single confocal slices of *E*12.5 control and *Tbx4-rtTA;tet-O-Cre;Srf*^*fl/fl*^ lungs immunostained for Ecad and Ki67. White dashed line indicates border of the epithelium. Arrowheads point to smooth muscle cells with high Ki67 intensity. (**H**) Quantification of nuclear Ki67 intensity in smooth muscle cells of mutants and controls compared by t-test (n = 3). (**I**) Percentages of control and *Tbx4-rtTA;tet-O-Cre;Srf*^*fl/fl*^ smooth muscle cells at each phase of the cell cycle based on scRNA-seq data. Scale bars indicate 50 μm.

To test the plausibility of this hypothesis, we compiled a list of genes that serve as markers for contractile and synthetic adult smooth muscle (Yap et al., 2021, Augstein et al., 2018, Wang et al., 2021, Jaslove and Nelson, 2018, Chettimada et al., 2016, Owens et al., 2004) and clustered *Srf*-mutant and control embryonic airway smooth muscle cells based on their expression of these genes (**Fig. 4B, Supplementary Table 1**). Contractile and synthetic markers tend to cluster together, as do mutant and control smooth muscle. Consistent with our hypothesis, contractile markers are strongly enriched in control smooth muscle, whereas synthetic markers are enriched in *Srf*-null smooth muscle. Based on marker expression, wildtype (mature) airway smooth muscle exhibits a classical contractile phenotype. In contrast, *Srf*-null (immature) smooth muscle displays a gene-expression program consistent with a classical synthetic phenotype.

We confirmed this observation using immunofluorescence and RNAscope analysis for markers of synthetic smooth muscle (Sox9 and Col21a1) and cell proliferation (Ki67). We observed dense accumulation of Sox9^+^ cells in the mesenchyme around the branching epithelium (measured between the first and second domain branches of the left lobe) of *Srf*-mutant lungs, but not controls (**Fig. 4C-D**). Additionally, the Sox9^+^ cell layer in mutants is significantly thicker than the αSMA^+^ cell layer in controls (**Fig. 4E**). *Col2a1* is present in the epithelium and in the cartilage precursors lining the lateral side of the upper primary bronchi in both controls and mutants (**Fig. 4F**), as expected given its known gene expression patterns (Andrikopoulos et al., 1992, Ng et al., 1997). Additionally, we observed *Col2a1* expression in *Acta2*^-^*Myocd*^*+*^ mutant smooth muscle cells but not *Acta2*^+^*Myocd*^*+*^ controls (**Fig. 4F**). Ki67 is enriched in the smooth muscle cells of the mutant compared to the control, suggesting that more *Srf*-null smooth muscle cells are actively proliferating (**Fig. 4G-H**). Consistently, cell-cycle scoring of our scRNA-seq data revealed that more cells of the control are in G1 phase and that more cells of the mutant are in S phase (**Fig. 4I**). Given that *Srf*-null smooth muscle expresses Sox9 and Col2a1 and that it is highly proliferative, we conclude that the loss of Srf arrests this population in a synthetic smooth muscle state. Our data thus reveal that *Srf* promotes the contractile smooth muscle phenotype in embryonic airway smooth muscle.

## Discussion

The airway epithelium branches normally in embryos lacking *Srf* in the pulmonary mesenchyme. Consistently, *Srf*-null mesenchyme displays normal patterning and passive mechanical properties during the early stages of lung development. Close inspection revealed that *Srf*-mutant lungs contain a population of smooth muscle cells that lacks classical markers of contractile smooth muscle but that is otherwise transcriptionally and functionally similar to control smooth muscle. This *Srf*-null population resembles immature smooth muscle and displays markers of the classical synthetic smooth muscle phenotype, suggesting that embryonic airway smooth muscle matures along a continuum from synthetic to contractile states. Phenotyping switching of smooth muscle in the adult might therefore represent a reactivation/reversal of normal embryonic programming.

Our findings demonstrate that SRF-independent aspects of airway smooth muscle differentiation are sufficient to support epithelial branching morphogenesis. Since these early stages are the important sculptors of the airway epithelium (and not the later stages of smooth muscle maturation) (Goodwin et al., 2019, Kim et al., 2015), it is unsurprising that SRF/myocardin-induced transcription is dispensable for branching morphogenesis of the airway epithelium. *Srf* (or *Myocd*) deletion leads to only a subtle change in epithelial morphology: a narrowing of branch stalks, despite the loss of the contractile smooth muscle phenotype (**Fig. 1D**) (Young et al., 2020). Our results provide an explanation for this seemingly contradictory observation. Accumulation of proliferative synthetic smooth muscle may squeeze the airways simply by wrapping the epithelium with a larger number of cells with elevated stiffness and mechanical tone, without the need for contractile smooth muscle machinery. This concept is illustrated by reconstructed cross-sections through the airways, which reveal that a dense layer of Sox9^+^ cells encircles the epithelium in the mutant, twice as thick as the layer of αSMA^+^ cells in the control (**Fig. 4C, E**). Further support for this model comes from our observation that the ability of the *Srf*-null mesenchyme to mechanically sculpt the epithelium into bifurcations (**Fig. 2N-O**) is the same as the control.

Overall, our findings suggest that synthetic smooth muscle is capable of providing its adjacent epithelium with some of the same mechanical signals as contractile smooth muscle. As a result, this phenotypic switch in smooth muscle has no consequence for branching morphogenesis in the embryonic mouse lung. Future studies are needed to determine whether a synthetic smooth muscle phenotype is conserved during the development of other embryonic organs.

## Abbreviations

αSMA: α-smooth muscle actin
Bmp4: bone morphogenetic protein 4
Ecad: E-cadherin
Lef1: lymphoid enhancer binding factor 1
pMLC: phosphorylated myosin light chain
SRF: serum response factor
Tbx3: T-box transcription factor 3

## Acknowledgements

We thank Dr. Wei Wang and the Genomics Core Facility of Princeton University for assistance with transcriptomics. We are grateful to Dr. Wei Shi (Keck School of Medicine of USC) for generously providing us with the *Tbx4-rtTA;tet-O-Cre* mouse line. We thank the Confocal Imaging Facility, a Nikon Center of Excellence, in the Department of Molecular Biology at Princeton University for instrument use and technical advice. We are also grateful to members of the Tissue Morphodynamics Group for helpful discussions and feedback on the manuscript. This work was supported by the NIH (HD099030, HL120142, and HL164861) and an HHMI Faculty Scholars Award to C.M.N. K.G. was supported in part by a postgraduate scholarship-doctoral (PGS-D) from the Natural Sciences and Engineering Research Council of Canada, the Dr. Margaret McWilliams Predoctoral Fellowship from the Canadian Federation of University Women, the Princeton University Procter Fellowship, and an American Heart Association Predoctoral Fellowship.

## Author contributions

K.G. and C.M.N. conceptualized the study, designed the experiments, interpreted the data, and wrote the manuscript. K.G., B.L., and A.B. performed the experiments and collected the data. All authors provided input on the final manuscript.

## STAR Methods

### RESOURCE AVAILABILITY

#### Lead Contact

- Further information and requests for resources and reagents should be directed to and will be fulfilled by the lead contact, Celeste M. Nelson.

#### Materials availability

- This study did not generate new unique reagents.

#### Data and code availability

- Single-cell RNA-seq data have been deposited at GEO and are publicly available as of the date of publication. Accession numbers are listed in the key resources table.
- Any additional information required to reanalyze the data reported in this paper is available from the lead contact upon request.

### EXPERIMENTAL MODEL AND SUBJECT DETAILS

#### Mice

Breeding of Srf-flox (Jackson Laboratory Stock 006658), Rosa26-mTmG (to generate *Srf-flox;mTmG* mice; Jackson Laboratory Stock 007676), and Tbx4-rtTA;tet-O-Cre (Zhang et al., 2013) (gift of Dr. Wei Shi) mice and isolation of embryos were carried out in accordance with institutional guidelines following the NIH Guide for the Care and Use of Laboratory Animals and approved by Princeton’s Institutional Animal Care and Use Committee. Doxycycline was administered in drinking water at 0.5 mg/ml starting at *E*5.5 or *E*6.5. Genotyping was carried out by isolating DNA from ear punches of pups or from the head of each embryo, followed by PCR and gel electrophoresis. In experiments using *Srf-flox;mTmG* mice, Cre and Tbx4 genotype was determined by the presence of GFP. The primer sequences for Cre were GCATTACCGGTCGATGCAACGAGTGATGAG and GAGTGAACGAACCTGGTCGAAATCAGTGCG, and the primer sequences for Tbx4-rtTA were GGAAGGCGAGTCATGGCAAGA and AGGTCAAAGTCGTCAAGGGCA. The primer sequences for Srf-flox and mTmG are all provided on the Jackson Laboratory website.

## METHOD DETAILS

### Wholemount immunofluorescence staining and imaging

Isolated lungs were fixed in 4% paraformaldehyde in PBS for 30 minutes at 4ºC. Lungs were washed four times for 15 minutes with PBST (0.1% Triton X-100 in PBS) and then blocked with 5% goat serum and 0.1% BSA for one hour. Samples were then incubated with primary antibodies against αSMA (Sigma a5228, 1:400 or Abcam ab5694, 1:200), E-cadherin (Cell Signaling 3195, 1:200 or Invitrogen 13-1900, 1:200), Lef1 (Cell Signaling 2230, 1:200), pMLC2 (Cell Signaling 3671, 1:200), RFP (Abcam ab62341, 1:400), Sox9 (Sigma AB5535, 1:1000), or Tbx3 (Invitrogen 42-4800, 1:200), followed by incubation with Alexa Fluor-conjugated secondary antibodies (1:400; Thermo Fisher Scientific A11007, A21244, A11012, A21240 and A11006) or Alexa Fluor-conjugated phalloidin (1:400, Thermo Fisher Scientific A12380 or A12379) and then counterstained with Hoechst (1:1000). Lungs were dehydrated in a methanol or isopropanol (for phalloidin-stained samples) series and cleared with Murray’s clear (1:2 ratio of benzyl alcohol to benzyl benzoate). Samples were imaged using a spinning disk confocal (BioVision X-Light V2) fitted to an inverted microscope.

### Fluorescence in situ hybridization

Isolated lungs were immediately placed in freshly prepared 4% paraformaldehyde in PBS for 24 hours at 4ºC. Lungs were then washed in PBS, followed by 20% sucrose and then 30% sucrose in PBS at 4ºC until they sank (less than 24 hours). All reagents were prepared using autoclaved and RNaseZAP-treated (Sigma R2020) water. Lungs were then embedded in OCT (Tissue Tek), frozen on dry ice, and stored at -80ºC. Sectioning was performed on a Leica CM3050S cryostat. Sample blocks were equilibrated at -20ºC in the cryostat for 1 hour and then sliced into 10-μm-thick sections onto Superfrost Plus slides (Fisherbrand). Fluorescence in situ hybridization was performed using the standard RNAScope Multiplex Fluorescent V2 Assay (ACD) protocol for fixed-frozen samples. Probes used were for *Mus musculus Myocd* (channel 1, 581051), *Col2a1* (channel 2, 407221), *Bmp4* (channel 2, 401301), and *Acta2* (channel 3, 319531). Fluorophores were Opal 520 and Opal 620 (Akoya Biosciences FP1487001KT and FP1495001KT). Sections were imaged on a Nikon A1RSi confocal microscope with a 20× objective.

### Organ culture and live imaging of contractions

Lung explants were cultured *ex vivo* following established protocols (Carraro et al., 2010). Lungs were dissected in cold PBS supplemented with antibiotics (50 units/ml of penicillin and streptomycin) and cultured on porous membranes (nucleopore polycarbonate track-etch membrane, 8 μm pore size, 25 mm diameter; Whatman) floating on DMEM/F12 medium (without HEPES) supplemented with 5% fetal bovine serum (FBS, heat inactivated; Atlanta Biologicals) and antibiotics (50 units/ml of penicillin and streptomycin) within a glass-bottom dish. Lungs were cultured within a stage-top incubator (Pathology Devices) on an inverted microscope (Nikon Ti) and imaged under brightfield (1-2 ms exposure). Every hour for 12 hours, a five-minute timelapse was taken with frames acquired every three seconds. Timelapses from hours 6 to 12 were analyzed for the presence of spontaneous smooth muscle contractions, visible by deformations of the epithelium, and compared using a two-sided t-test in GraphPad Prism.

### scRNA-seq experiments

Lungs isolated at *E*12.5 from two litters of *Tbx4-rtTA;tet-O-Cre;Srf*^*fl/fl*^ mutants (7 lungs) and *Srf*^*fl/fl*^ littermate controls (6 lungs) were dissected in sterile-filtered PBS, mechanically dissociated using fine tungsten needles (Fine Science Tools), and then further dissociated in dispase (Corning 354235) on ice for 15 minutes to create single-cell suspensions. Dispase was inactivated using DMEM/F12 medium (without HEPES) with 5% FBS and antibiotics (50 units/ml of penicillin and streptomycin) and the cell suspensions were passed through 40-μm-diameter mesh filters. scRNA-seq library preparation and sequencing were carried out by the Princeton Genomics Core Facility using the Chromium Single Cell 3’ Library and Gel Bead Kit v3 on the Chromium Controller (10X Genomics) following manufacturer protocols. Illumina sequencing libraries were prepared from the PCR-amplified cDNA from mutant lungs and controls using the Nextra DNA library prep kit (Illumina). Libraries were sequenced on a NovaSeq 6000 S Prime flowcell (Illumina) as paired-end 28 + 94 nucleotide reads, following manufacturer protocols. Base calling was performed and raw sequencing reads were filtered using the Illumina sequencer control software. The 10x CellRanger software version 3.0.2 was used to run the count pipeline with default settings on all FASTQ files from each sample to generate gene-barcode matrices using the *Mus musculus* reference genome mm10-1.2.0. Data are available from GEO (GSE203171).

## QUANTIFICATION AND STATISTICAL ANALYSES

### scRNA-seq analysis

scRNA-seq data were imported into R and processed using the Seurat package following the recommended pipeline (Butler et al., 2018). Clusters and expression levels of genes of interest were visualized using the uniform manifold approximation and projection (UMAP) dimensional reduction. KEGG enrichment analysis was performed on markers for smooth muscle clusters from mutants and controls using the clusterProfiler package in R (Yu et al., 2012). Diffusion analyses were carried out using the Destiny package in R (Haghverdi et al., 2015). First, we computationally isolated mesenchymal cells from the control dataset and merged them with the smooth muscle cells of the mutant dataset. The resultant Seurat object was passed through the FindVariableFeatures and ScaleData functions. Diffusion analysis was performed on only the control cells, followed by projection of the mutant cells onto the existing diffusion map.

### Image and statistical analyses

Quantifications of airway diameter, projected epithelial and whole lung areas, and number of terminal branches were made manually in Image J and compared using a two-sided t-test in GraphPad Prism. For immunofluorescence analysis, intensity profiles were measured either around the airways and plotted in polar coordinates or along lines emanating from the epithelium using simple MATLAB pipelines. Images were first background subtracted and then average pixel intensities in 16 by 16 pixel (5.76 by 5.76 μm) windows were measured along contours or lines. For intensities measured around the airway epithelium, we compared the mean intensity along the contour of mutants and controls using a two-sided t-test in MATLAB. For intensities measured along a line emanating from the epithelium, we compared mutants and controls by two-way ANOVA in GraphPad Prism. For Ki67 immunofluorescence analysis, we measured the nuclear intensity of Ki67 in the mesenchyme adjacent to airway stalks. Background intensity was subtracted, and 20 nuclei were measured per lung.

## Supplementary Figures

**Supplementary Figure 1,related to Figure 1.**
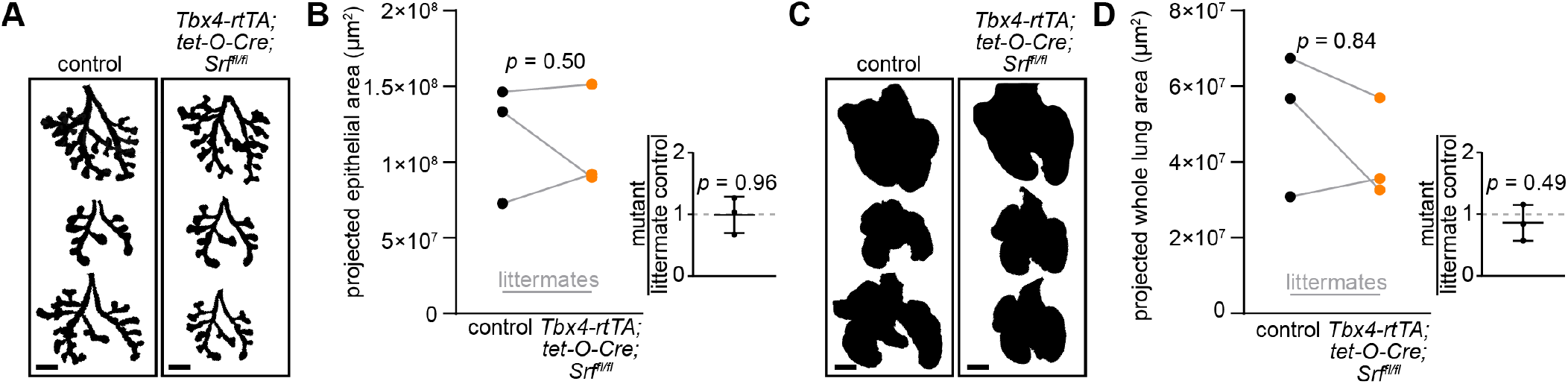
Lung size is unaltered in *Srf* mutants. (**A**) Thresholded images of z-projections of the airway epithelium (based on Ecad immunofluorescence). Scale bars indicate 200 μm. (**B**) Projected epithelial area in *E*12.5 *Tbx4-rtTA;tet-O-Cre;Srf*^*fl/fl*^ and littermate controls, connected by gray lines and compared using a t-test. Inset shows projected epithelial area in mutant lungs divided by that of littermate controls compared to a hypothetical mean of 1 using a one sample t-test. Error bars show s.d. (**C**) Thresholded images of z-projections of the entire lung (based on αSMA background immunofluorescence). Scale bars indicate 200 μm. (**D**) Projected whole lung area in *E*12.5 *Tbx4-rtTA;tet-O-Cre;Srf*^*fl/fl*^ and littermate controls, connected by gray lines and compared using a t-test. Inset shows projected area of whole lungs in mutants divided by that of littermate controls compared to a hypothetical mean of 1 using a one sample t-test. Error bars show s.d.

**Supplementary Figure 2,related to Figure 2.**
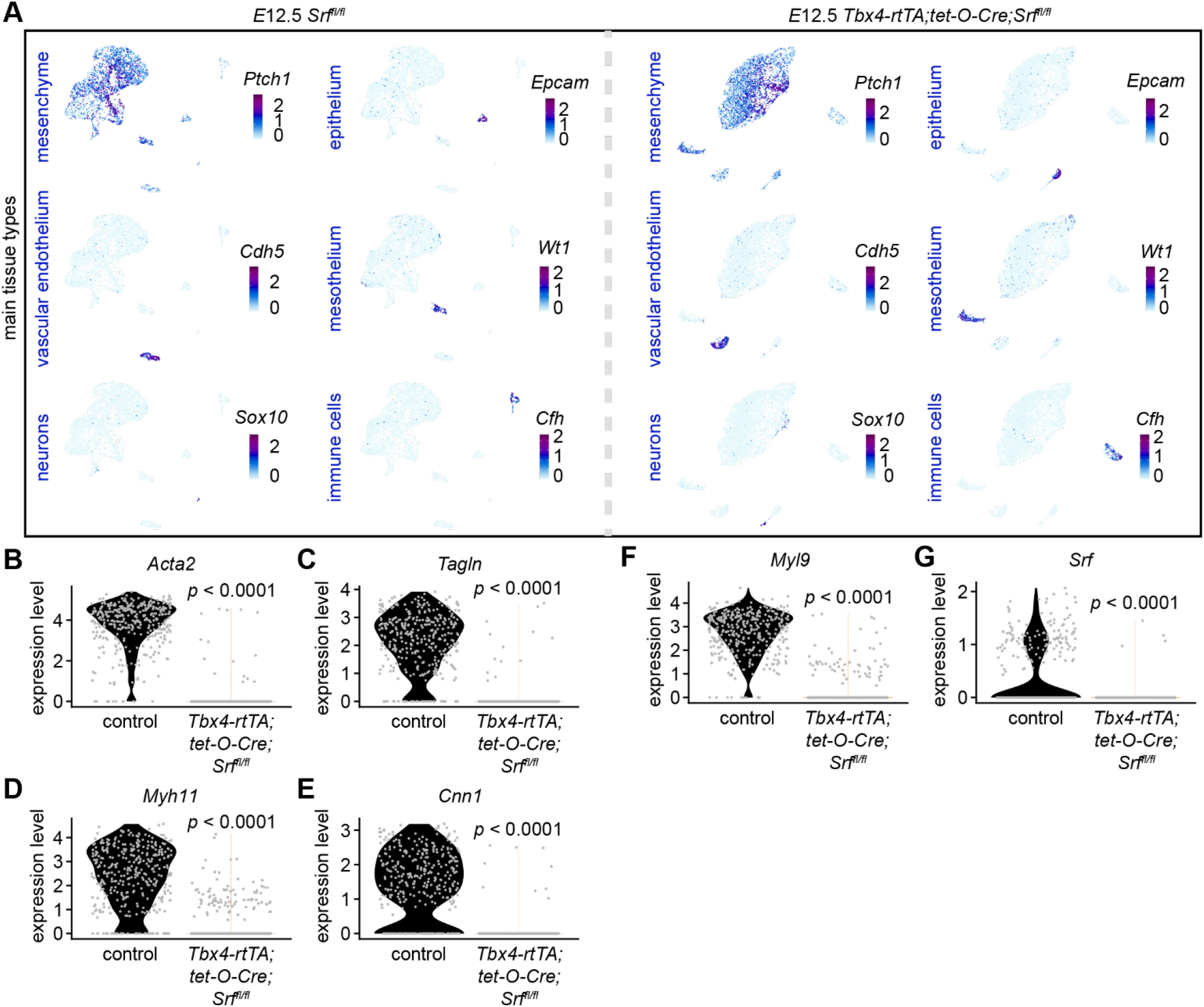
scRNA-seq of control and *Srf*-mutant lungs. (**A**) UMAPs of cells isolated from *E*12.5 *Tbx4-rtTA;tet-O-Cre;Srf*^*fl/fl*^ and littermate control lungs color-coded according to the expression levels of marker genes for the major cell types within the embryonic lung. (**B-G**) Expression levels of canonical markers for the contractile smooth muscle phenotype in smooth muscle cells from control and *Tbx4-rtTA;tet-O-Cre;Srf*^*fl/fl*^ lungs.

**Supplementary Figure 3,related to Figure 3.**
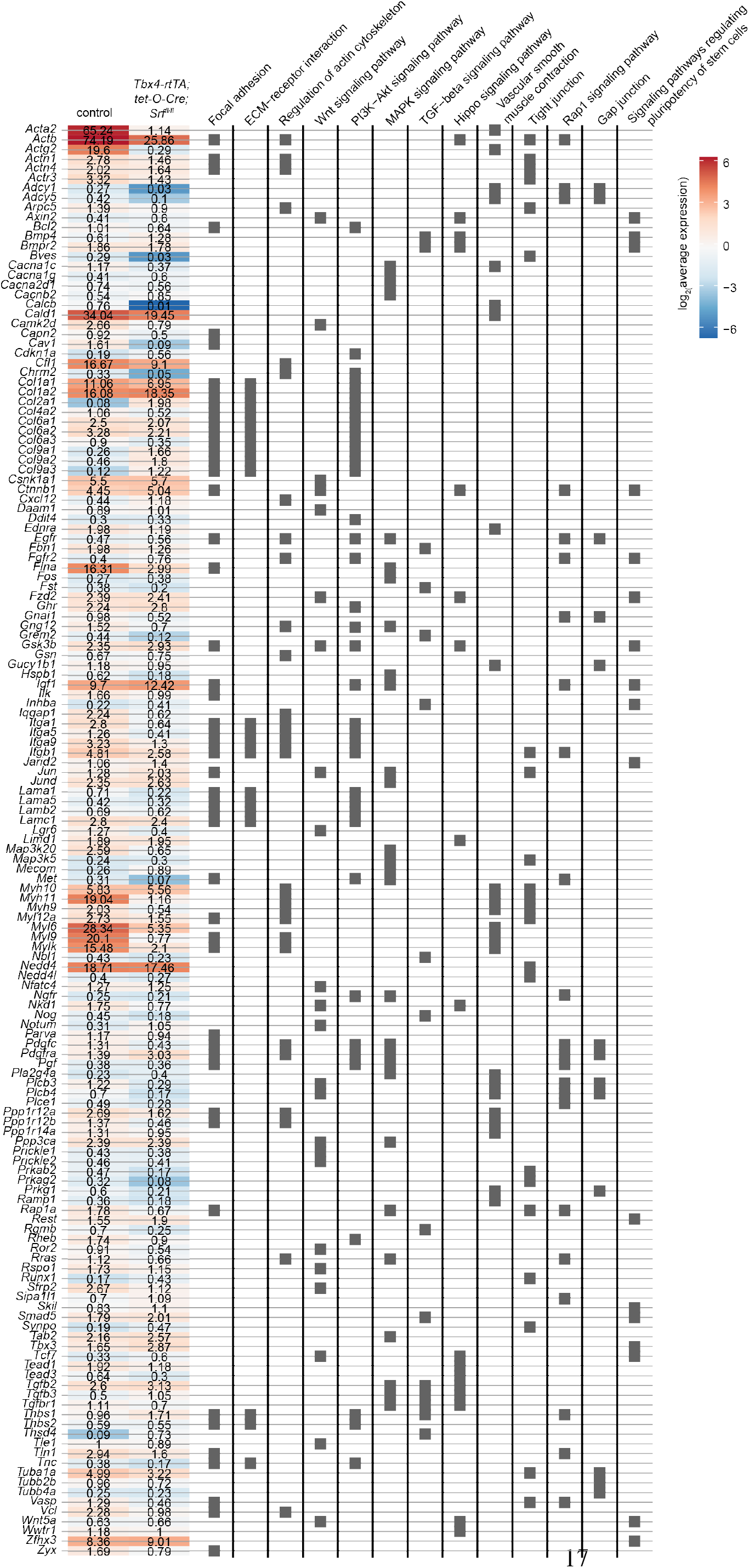
Average expression of genes from enriched KEGG terms. Average expression of genes in each KEGG term from **Fig. 3F**. Color scale shows log2 of average gene expression, and the scaled expression level of each gene is indicated.

**Supplementary Table 1.**
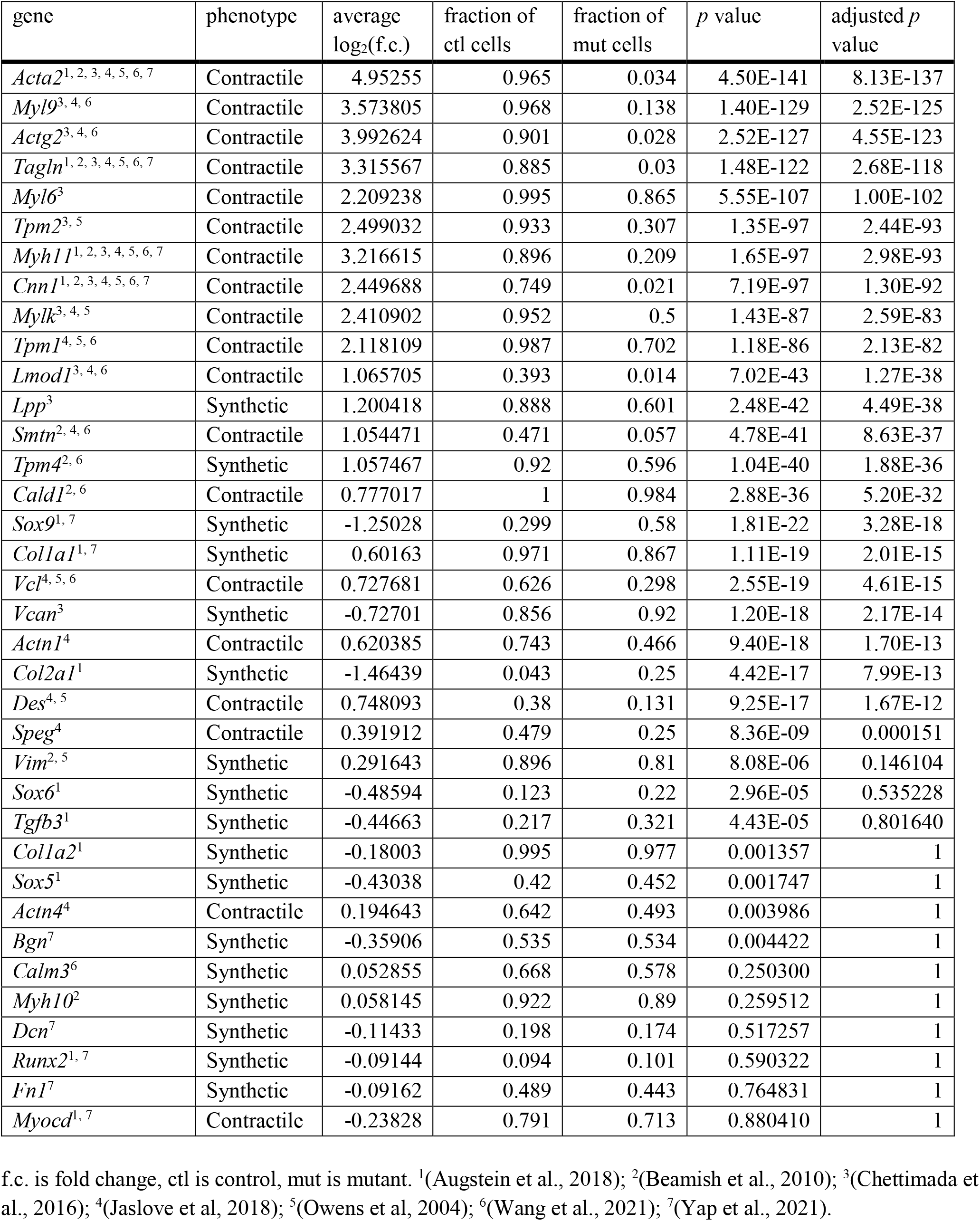
Expression of contractile and synthetic smooth muscle markers in *Srf*-mutant lungs and controls.

## References

Andrikopoulos, K., Suzuki, H. R., Solursh, M. & Ramirez, F. 1992. Localization of pro-alpha 2(V) collagen transcripts in the tissues of the developing mouse embryo. Dev Dyn, 195, 113–20.

Arsenian, S., Weinhold, B., Oelgeschlager, M., Ruther, U. & Nordheim, A. 1998. Serum response factor is essential for mesoderm formation during mouse embryogenesis. EMBO J, 17, 6289–99.

Augstein, A., Mierke, J., Poitz, D. M. & Strasser, R. H. 2018. Sox9 is increased in arterial plaque and stenosis, associated with synthetic phenotype of vascular smooth muscle cells and causes alterations in extracellular matrix and calcification. Biochim Biophys Acta Mol Basis Dis, 1864, 2526–2537.

Boucherat, O., Landry-Truchon, K., Berube-Simard, F. A., Houde, N., Beuret, L., Lezmi, G., Foulkes, W. D., Delacourt, C., Charron, J. & Jeannotte, L. 2015. Epithelial inactivation of Yy1 abrogates lung branching morphogenesis. Development, 142, 2981–95.

Butler, A., Hoffman, P., Smibert, P., Papalexi, E. & Satija, R. 2018. Integrating single-cell transcriptomic data across different conditions, technologies, and species. Nat Biotechnol, 36, 411–420.

Carraro, G., Del Moral, P. M. & Warburton, D. 2010. Mouse embryonic lung culture, a system to evaluate the molecular mechanisms of branching. J Vis Exp.

Chettimada, S., Joshi, S. R., Dhagia, V., Aiezza, A., 2nd, Lincoln, T. M., Gupte, R., Miano, J. M. & Gupte, S. A. 2016. Vascular smooth muscle cell contractile protein expression is increased through protein kinase G-dependent and -independent pathways by glucose-6-phosphate dehydrogenase inhibition and deficiency. Am J Physiol Heart Circ Physiol, 311, H904–H912.

Donadon, M. & Santoro, M. M. 2021. The origin and mechanisms of smooth muscle cell development in vertebrates. Development, 148.

Du, K. L., Ip, H. S., Li, J., Chen, M., Dandre, F., Yu, W., Lu, M. M., Owens, G. K. & Parmacek, M. S. 2003. Myocardin is a critical serum response factor cofactor in the transcriptional program regulating smooth muscle cell differentiation. Mol Cell Biol, 23, 2425–37.

Feng, H. Z., Wang, H., Takahashi, K. & Jin, J. P. 2019. Double deletion of calponin 1 and calponin 2 in mice decreases systemic blood pressure with blunted length-tension response of aortic smooth muscle. J Mol Cell Cardiol, 129, 49–57.

Goodwin, K., Jaslove, J. M., Tao, H., Zhu, M., Hopyan, S. & Nelson, C. M. 2022. Patterning the embryonic pulmonary mesenchyme. iScience, 25, 103838.

Goodwin, K., Mao, S., Guyomar, T., Miller, E., Radisky, D. C., Kosmrlj, A. & Nelson, C. M. 2019. Smooth muscle differentiation shapes domain branches during mouse lung development. Development, 146.

Gunst, S. J. & Zhang, W. 2008. Actin cytoskeletal dynamics in smooth muscle: a new paradigm for the regulation of smooth muscle contraction. Am J Physiol Cell Physiol, 295, C576–87.

Haghverdi, L., Buettner, F. & Theis, F. J. 2015. Diffusion maps for high-dimensional single-cell analysis of differentiation data. Bioinformatics, 31, 2989–98.

He, H., Huang, M., Sun, S., Wu, Y. & Lin, X. 2017. Epithelial heparan sulfate regulates Sonic Hedgehog signaling in lung development. PLoS Genet, 13, e1006992.

Hoofnagle, M. H., Neppl, R. L., Berzin, E. L., Teg Pipes, G. C., Olson, E. N., Wamhoff, B. W., Somlyo, A. V. & Owens, G. K. 2011. Myocardin is differentially required for the development of smooth muscle cells and cardiomyocytes. Am J Physiol Heart Circ Physiol, 300, H1707–21.

Jaslove, J. M., Goodwin, K., Sundarakrishnan, A., Spurlin, J. W., Mao, S., Kosmrlj, A. & Nelson, C. M. 2022. Transmural pressure signals through retinoic acid to regulate lung branching. Development, 149.

Jaslove, J. M. & Nelson, C. M. 2018. Smooth muscle: a stiff sculptor of epithelial shapes. Philos Trans R Soc Lond B Biol Sci, 373.

Kim, H. Y., Pang, M. F., Varner, V. D., Kojima, L., Miller, E., Radisky, D. C. & Nelson, C. M. 2015. Localized Smooth Muscle Differentiation Is Essential for Epithelial Bifurcation during Branching Morphogenesis of the Mammalian Lung. Dev Cell, 34, 719–26.

Li, S., Wang, D. Z., Wang, Z., Richardson, J. A. & Olson, E. N. 2003. The serum response factor coactivator myocardin is required for vascular smooth muscle development. Proc Natl Acad Sci U S A, 100, 9366–70.

Luo, L., Wang, L., Pare, P. D., Seow, C. Y. & Chitano, P. 2019. The Huxley crossbridge model as the basic mechanism for airway smooth muscle contraction. Am J Physiol Lung Cell Mol Physiol, 317, L235–L246.

Miano, J. M. 2015. Myocardin in biology and disease. J Biomed Res, 29, 3–19.

Ng, L. J., Wheatley, S., Muscat, G. E., Conway-Campbell, J., Bowles, J., Wright, E., Bell, D. M., Tam, P. P., Cheah, K. S. & Koopman, P. 1997. SOX9 binds DNA, activates transcription, and coexpresses with type II collagen during chondrogenesis in the mouse. Dev Biol, 183, 108–21.

Owens, G. K., Kumar, M. S. & Wamhoff, B. R. 2004. Molecular regulation of vascular smooth muscle cell differentiation in development and disease. Physiol Rev, 84, 767–801.

Rensen, S. S., Doevendans, P. A. & Van Eys, G. J. 2007. Regulation and characteristics of vascular smooth muscle cell phenotypic diversity. Neth Heart J, 15, 100–8.

Schildmeyer, L. A., Braun, R., Taffet, G., Debiasi, M., Burns, A. E., Bradley, A. & Schwartz, R. J. 2000. Impaired vascular contractility and blood pressure homeostasis in the smooth muscle alpha-actin null mouse. FASEB J, 14, 2213–20.

Sieck, G. C., Dogan, M., Young-Soo, H., Osorio Valencia, S. & Delmotte, P. 2019. Mechanisms underlying TNFalpha-induced enhancement of force generation in airway smooth muscle. Physiol Rep, 7, e14220.

Wang, G., Jacquet, L., Karamariti, E. & Xu, Q. 2015. Origin and differentiation of vascular smooth muscle cells. J Physiol, 593, 3013–30.

Wang, L., Rice, M., Swist, S., Kubin, T., Wu, F., Wang, S., Kraut, S., Weissmann, N., Bottger, T., Wheeler, M., Schneider, A. & Braun, T. 2021. BMP9 and BMP10 Act Directly on Vascular Smooth Muscle Cells for Generation and Maintenance of the Contractile State. Circulation, 143, 1394–1410.

Wang, Z., Wang, D. Z., Pipes, G. C. & Olson, E. N. 2003. Myocardin is a master regulator of smooth muscle gene expression. Proc Natl Acad Sci U S A, 100, 7129–34.

Yap, C., Mieremet, A., De Vries, C. J. M., Micha, D. & De Waard, V. 2021. Six Shades of Vascular Smooth Muscle Cells Illuminated by KLF4 (Kruppel-Like Factor 4). Arterioscler Thromb Vasc Biol, 41, 2693–2707.

Yi, L., Domyan, E. T., Lewandoski, M. & Sun, X. 2009. Fibroblast growth factor 9 signaling inhibits airway smooth muscle differentiation in mouse lung. Dev Dyn, 238, 123–37.

Young, R. E., Jones, M. K., Hines, E. A., Li, R., Luo, Y., Shi, W., Verheyden, J. M. & Sun, X. 2020. Smooth Muscle Differentiation Is Essential for Airway Size, Tracheal Cartilage Segmentation, but Dispensable for Epithelial Branching. Dev Cell, 53, 73–85 e5.

Yu, G., Wang, L. G., Han, Y. & He, Q. Y. 2012. clusterProfiler: an R package for comparing biological themes among gene clusters. OMICS, 16, 284–7.

Zhang, J. C., Kim, S., Helmke, B. P., Yu, W. W., Du, K. L., Lu, M. M., Strobeck, M., Yu, Q. & Parmacek, M. S. 2001. Analysis of SM22alpha-deficient mice reveals unanticipated insights into smooth muscle cell differentiation and function. Mol Cell Biol, 21, 1336–44.

Zhang, W., Menke, D. B., Jiang, M., Chen, H., Warburton, D., Turcatel, G., Lu, C. H., Xu, W., Luo, Y. & Shi, W. 2013. Spatial-temporal targeting of lung-specific mesenchyme by a Tbx4 enhancer. BMC Biol, 11, 111.

